# The SARS-CoV-2 Spike is a virulence determinant and plays a major role on the attenuated phenotype of Omicron virus in a feline model of infection

**DOI:** 10.1101/2023.10.09.561473

**Authors:** Mathias Martins, Mohammed Nooruzzaman, Jessie Lee Cunningham, Chengjin Ye, Leonardo Cardia Caserta, Nathaniel Jackson, Luis Martinez-Sobrido, Ying Fang, Diego G. Diel

**Author notes:** Authors contributed equally to the work (Mathias Martins, Mohammed Nooruzzaman, Jessie Lee Cunningham). Address correspondence to Diego G. Diel,.

## Abstract

To assess the role of the Omicron BA.1 Spike (S) protein in the pathogenesis of the severe acute respiratory coronavirus 2 (SARS-CoV-2), we generated recombinant viruses harboring the S D614G mutation (rWA1-D614G) and the Omicron BA.1 S gene (rWA1-Omi-S) in the backbone of the ancestral SARS-CoV-2 WA1 strain genome. The recombinant viruses were characterized *in vitro* and *in vivo*. Viral entry, cell-cell fusion, viral plaque size, and viral replication kinetics of the rWA1-Omi-S virus were markedly impaired when compared to the rWA1-D614G virus, demonstrating a lower fusogenicity and ability to spread cell-to-cell of rWA1-Omi-S. To assess the contribution of the Omicron BA.1 S protein to SARS-CoV-2 pathogenesis the pathogenicity of rWA1-D614G and rWA1-Omi-S viruses were compared using a feline model of infection. While the rWA1-D614G-inoculated cats became lethargic and showed increased body temperatures on days 2 and 3 post-infection (pi), rWA1-Omi-S-inoculated cats remained subclinical and gained weight throughout the 14-day experimental period. Animals inoculated with rWA1-D614G presented higher levels of infectious virus shedding in nasal secretions, when compared to rWA1-Omi-S-inoculated animals. In addition, tissue replication of the rWA1-Omi-S was markedly reduced compared to the rWA1-D614G, as evidenced by lower *in situ* viral RNA and lower viral load in tissues on days 3 and 5 pi. Histologic examination of the nasal turbinate and lungs revealed intense inflammatory infiltration in rWA1-D614G-inoculated animals, whereas rWA1-Omi-S-inoculated cats presented only mild to modest inflammation. Together, these results demonstrate that the S protein is a major virulence determinant for SARS-CoV-2 playing a major role for the attenuated phenotype of the Omicron virus.

**Author summary:** The SARS-CoV-2 Omicron sublineage BA.1 spread rapidly across the globe in late 2021/early 2022. Experimental studies have shown an overall lower pathogenicity of Omicron BA.1 when compared to the ancestral SARS-CoV-2 lineage B.1 (D614G). Recently, we have demonstrated that the Omicron BA.1.1 variant presents lower pathogenicity when compared to D614G (B.1) lineage in a feline model of SARS-CoV-2 infection. There are over 50 mutations in the Omicron genome, of which more than two thirds are present in the S gene. To assess the role of the Omicron BA.1 S on virus pathogenesis, recombinant viruses harboring the S D614G mutation (rWA1-D614G) and the Omicron BA.1 Spike gene (rWA1-Omi-S) in the backbone of the ancestral SARS-CoV-2 WA1 were characterized *in vitro* and *in vivo*. While the Omicron BA.1 S gene results in early entry into cells, the rWA1-Omi-S presents impaired cell-cell spread and fusogenic activity. Inoculation of cats with the recombinant viruses revealed an attenuated phenotype of rWA1-Omi-S, demonstrating a critical role for S protein on the pathogenicity of SARS-CoV-2 and indicating that the Omi-S is a major determinant of the attenuated disease phenotype of Omicron strains.

## Introduction

Severe acute respiratory syndrome coronavirus 2 (SARS-CoV-2) was first detected in a cluster of people presenting with severe pneumonia in Wuhan, Hubei province in China [1]. Current evidence on the viral origin suggest that horseshoe bats are the likely reservoir for the virus, with human spillover potentially involving a yet unidentified intermediate animal host in the Huanan Seafood Wholesale market in Wuhan. Notably, 20 of the first 47 human cases in Wuhan had an epidemiological link with the Huanan market where several live wild animal species, including many that are now known to be susceptible to SARS-CoV-2, were sold [2–4].

Since the emergence of SARS-CoV-2 in the human population, there have been 764 million confirmed coronavirus disease 2019 (COVID-19) cases and over 6.9 million deaths, reported to World Health Organization (WHO). While circulating in the human population, SARS-CoV-2 has continued to evolve, with new variants causing significant waves of infection worldwide. Importantly, several of the emerging variants present altered transmissibility, immune evasion capability, and pathogenicity [5]. The first mutation established on the SARS-CoV-2 genome was the D614G substitution in the Spike (S) protein, detected for the first time in February 2020 and which is present in all SARS-CoV-2 lineages circulating worldwide since then [6]. As genome mutations accumulated and viral properties changed over time, some SARS-CoV-2 lineages were classified as variants of concern (VOC) by the WHO [7]. As of October 2023, five VOC have been defined: Alpha, Beta, Gamma, Delta, and Omicron. All these VOC emerged independently and rapidly became dominant, outcompeting previous variants regionally or globally. The SARS-CoV-2 Omicron VOC (BA.1), emerged in late 2021, and rapidly achieved global predominance by early 2022 [8,9]. A marked increase in the cases numbers was observed worldwide when the Omicron BA.1 variant became predominant owing to high re-infection and vaccine breakthrough rates of this virus due to its ability to evade neutralizing antibody responses [10–15]. Hospitalizations and deaths, however, were lower during the Omicron BA.1 surge suggesting lower pathogenicity of the virus compared to previous SARS-CoV-2 lineages [16].

The Omicron BA.1 sublineage differs from the ancestral Wuhan-Hu-1 virus by 59 amino acids, with 37 of these changes present in the S protein, a key viral protein involved in virus entry (receptor engagement and fusion) [17]. Among these, 15 amino acid substitutions are in the receptor binding domain (RBD) with several additional mutations located near the furin cleavage site at the S1/S2 junction and near the S2ʹ site which could modulate host protease cleavage by furin and transmembrane serine proteases (TMPRSS2), respectively, potentially resulting in altered entry preference and fusogenicity, and likely affecting replication and pathogenicity [17–20]. The Omicron BA.1 is less dependent on TMPRSS2, less efficient in S cleavage, less fusogenic, and presents an altered propensity to utilize the plasma membrane and endosomal pathways for virus entry. Specific mutations in the S protein have been linked to the attenuated phenotype of the Omicron BA.1 variant [18,21]. Additionally, amino acid mutations in the receptor binding motif (RBM) contribute to escape from vaccine-induced humoral immune responses [17].

Experimental studies in animal models showed that the Omicron variant causes attenuated disease in humanized mice and hamsters [17,22,23]. Using a feline model - a natural susceptible animal species - we have demonstrated that the Omicron BA.1.1 variant presents lower pathogenicity when compared to D614G (B.1) and the Delta lineages [24]. To assess the contribution of the S gene in this viral phenotype, we generated recombinant viruses based on the ancestral SARS-CoV-2 WA-1 strain backbone harboring the S D614G mutation (rWA1-D614G) and the Omicron BA.1 S gene (rWA1-Omi-S). The role of the S gene on virus entry/infectivity, fusion and replication were characterized *in vitro*, and the infection dynamics, tissue tropism, and pathogenicity of the viruses were assessed and compared *in vivo* using the highly susceptible feline model of infection and pathogenesis.

## Results

### The Omicron BA.1 S induces efficient viral entry, but leads to reduced replication of SARS-CoV-2 *in vitro*

The role and contribution of Omicron BA.1 S for virus infectivity, fusogenicity, cell-to-cell spread, and replication were investigated in cell culture *in vitro*. Two recombinant viruses based on the ancestral SARS-CoV-2 WA-1 genome backbone harboring the S D614G mutation (rWA1-D614G) and the Omicron BA.1 S gene (rWA1-Omi-S) were generated and rescued (Fig 1A). Initially, infectivity of rWA1-D614G and rWA1-Omi-S were investigated in cells stably expressing human (h) or cat (c) angiotensin converting enzyme 2 (hACE2_BHK21 and cACE2_BHK21, respectively), Vero E6, and Vero E6 TMPRSS2 cells and virus infectivity was determined by immunofluorescence assay (IFA) followed by quantification of infected cells. In both ACE2 expressing BHK21 cells and in Vero E6 TMPRSS2 cells, the rWA1-Omi-S virus showed higher infectivity at early time points of 4- and 8-hours post-infection (hpi) when compared to the rWA1-D614G (Figs 1B and 1C). At 12 hpi, both viruses showed comparable infectivity in hACE2_BHK21 and cACE2_BHK21 cells as evidenced by similar percentage of infected cells; however, from 12 hpi onwards the rWA1-D614G overtook the rWA1-Omi-S virus and at 24 hpi a significantly higher percentage of infected cells were noted in rWA1-D614G infected cells (Figs 1B and 1C). On the other hand, in Vero E6 cells, both recombinant viruses showed comparable infectivity throughout the course of infection. We then performed multi-step growth curves with both rWA1-D614G and rWA1-Omi-S in hACE2 and cACE2 expressing BHK21 cells, and Vero E6 cells, to assess the effect of S on virus replication. In both ACE2 expressing BHK21 cells, the rWA1-Omi-S virus presented slightly lower viral yields, suggesting lower replication ability of this virus, in comparison to the ancestral rWA1-D614G virus (Fig 1D). Whereas, both recombinant viruses presented comparable replication kinetics in Vero E6 cells (Fig 1D). These results are consistent with the infectivity assays performed in these cells (Figs 1B and 1C).

**FIG 1.**
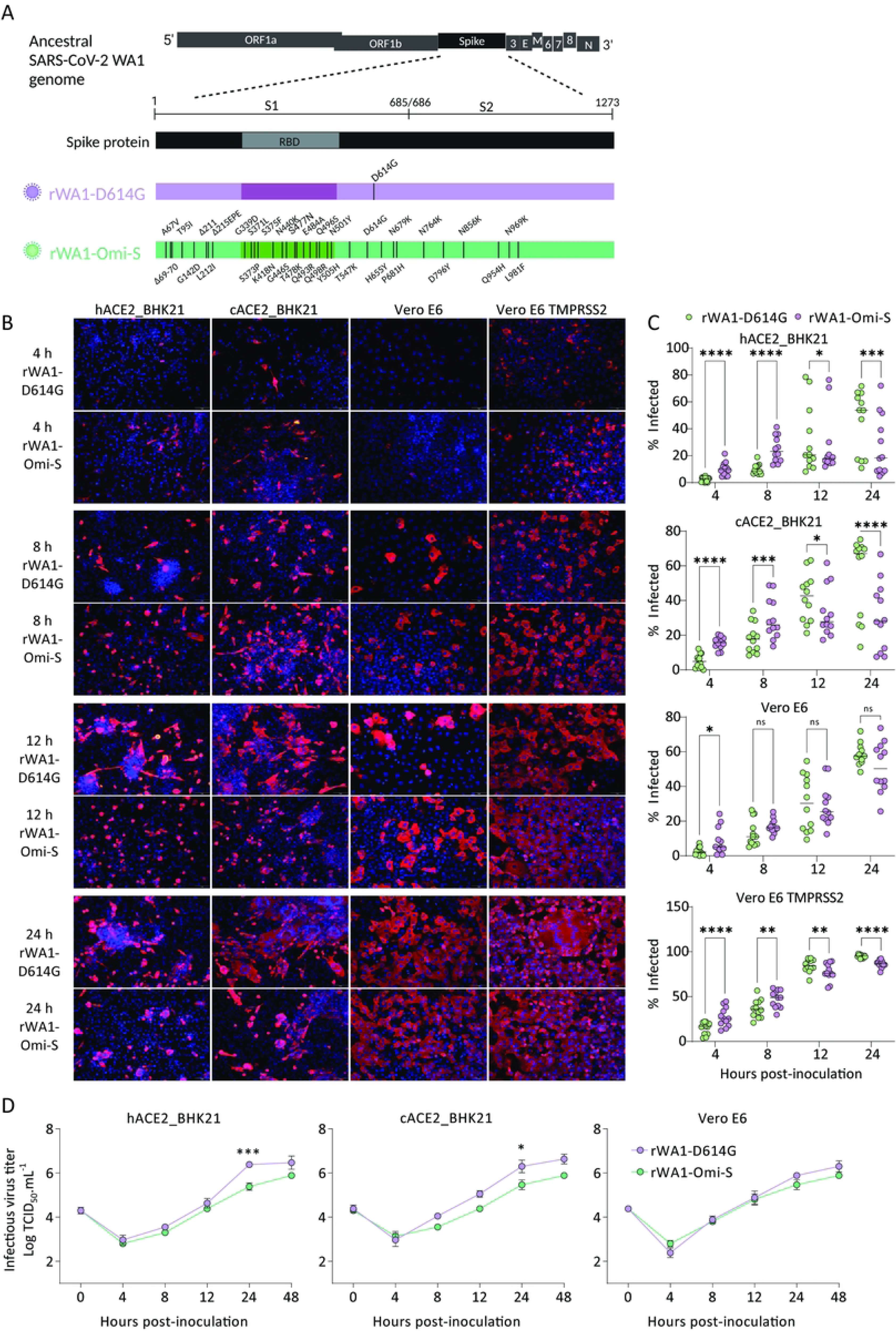
The role and contribution of Omicron BA.1 S to SARS-CoV-2 infectivity and replication. (A) Schematic presentation of the rSARS-CoV-2 harboring D614G (rWA1-D614G) and Omicron BA.1 S (rWA1-Omi-S). (B) IFA. BHK21 stably expressing human (hACE2_BHK21), and cat (cACE2_BHK21) ACE2, Vero E6, and Vero E6 TMPRSS2 cells were infected (MOI = 0.1) with rWA1-D614G and rWA1-Omi-S viruses and incubated at 37°C. At the indicated time points post-infection, cells were fixed and stained with a SARS-CoV-2 antibody against the viral nucleocapsid (N) protein (red) and DAPI (nucleus, blue). (C) Cells infected in B were counted and infectivity expressed as the percentage of infected cells based on the number of N positive cells over the total number of cells. A total of four fields were counted per experiment. Three independent experiments were performed. (D) Multi-step growth curves. hACE2_BHK21, cACE2_BHK21, and Vero E6 cells were infected (MOI = 0.1) with rWA1-D614G and rWA1-Omi-S and incubated at 37°C. Cells were harvested at the indicated times post-infection, and virus titers were determined by limiting dilutions and expressed as TCID_50_ per ml. The limit of detection (LOD) for infectious virus titration was 10^1.05^ TCID_50_.mL^-1^. Data are the means of three independent experiments (*n = 3*) ± SEM. 2-way ANOVA followed by multiple comparisons test, * *p* < 0.05, ** *p* < 0.01, *** *p* < 0.001 and **** *p* < 0.0001.

### The Omicron BA.1 S reduces fusogenicity and cell-to-cell spread of SARS-CoV-2 *in vitro*

The fusogenic activity of the Omicron BA.1 was evaluated and compared to the D614G S. For this, BHK21 cells stably expressing hACE2 or cACE2 were transfected with plasmids encoding the S gene from D614G and Omicron BA.1 and the fusogenic activity of the respective variant proteins were evaluated by determining the number of nuclei incorporated into the syncytia observed in transfected cells. Expression of Omicron BA.1 S in both hACE2 and cACE2-expressing BHK21 cells resulted in lower fusogenic activity as evidenced by significantly (*p* < 0.001) lower syncytia formation/nuclei incorporation when compared to expression of the ancestral D614G S protein (Figs 2A and 2B). Notably, the lower fusogenicity of Omicron BA.1 S protein was confirmed in the context of virus infection. BHK21 cells expressing hACE2 or cACE2 infected with rWA1-Omi-S presented markedly lower fusogenicity as evidenced by lower number and smaller syncytia when compared to cells infected with the ancestral rWA1-D614G virus (Figs 2C and 2D). Next, we assessed the effect of Omicron S protein on cell-to-cell spread of SARS-CoV-2 by using plaque assays. The rWA1-Omi-S formed significantly smaller plaques (*p* < 0.001) when compared to the rWA1-D614G virus in all three cell lines tested (hACE2_BHK21, cACE2_BHK21 and Vero E6) (Figs 2E and 2F), indicating a reduced ability of the rWA1-Omi-S to spread from cell-to-cell and confirming the contribution of S protein for this phenotype. Collectively, these results demonstrate that while the Omicron BA.1 S enables efficient virus entry and infectivity, it reduces the overall fusogenicity and ability of the rWA-Omi-S virus to spread from cell-to-cell.

**FIG 2.**
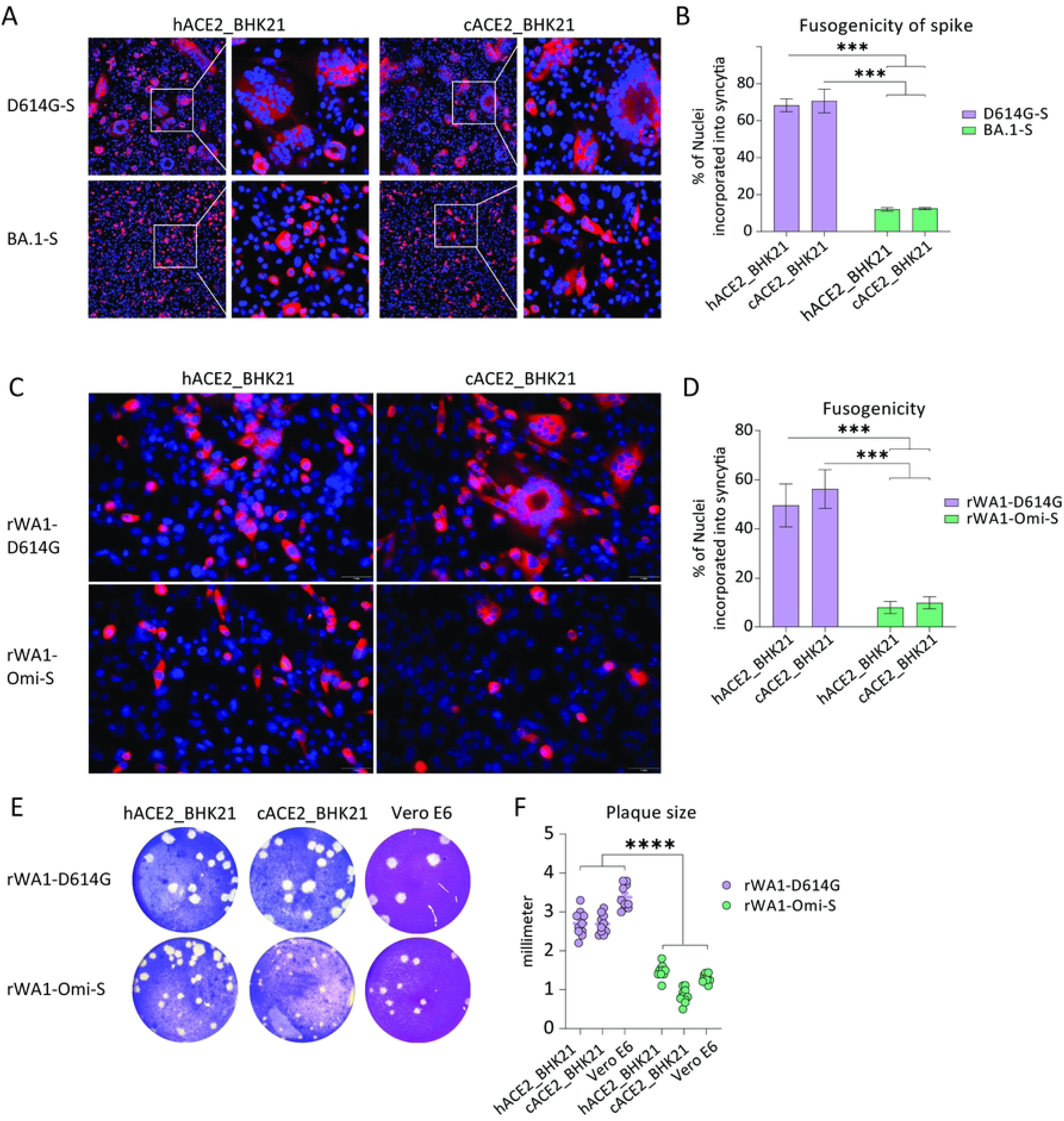
The rSARS-CoV-2 WA-1 harboring the Omicron BA.1 S gene (rWA1-Omi-S) presents reduced fusogenicity *in vitro*. (A) BHK21 cells stably expressing human (hACE2_BHK21) or cat ACE2 (cACE2_BHK21) were transfected with plasmids carrying either the D614G or the Omicron BA.1 S proteins and syncytia formation was visualized by IFA. (B) The fusogenic activity of the D614G and the Omicron BA.1 S was quantified by counting the incorporation of nuclei into syncytia observed in A. The percentage of nuclei observed per field (*n* = 5) and incorporated into syncytia were calculated over the total number of nuclei per field. (C) Cells stably expressing human (hACE2_BHK21) and cat (cACE2_BHK21) ACE2 were infected (MOI 0.1) with rWA1-D614G and rWA1-Omi-S and incubated for 24 h at 37°C. Syncytia formation was visualized by IFA using an N-specific mAb. (D) The fusogenic activity of the rWA-D614G and the rWA1-Omi-S were quantified by counting the incorporation of nuclei into syncytia observed in C. The percentage of nuclei observed per field (*n* = 4) and incorporated into syncytia were calculated over the total number of nuclei per field. (E) Plaque phenotype of the rWA1-D614G and rWA1-Omi-S viruses. hACE2_BHK21, cACE2_BHK21, and Vero E6 cells were infected with rWA1-D614G and rWA1-Omi-S and overlaid with 0.5% agarose medium. Plates were incubated at 37°C for 72 h, the agar overlay was removed, cells were fixed, and the monolayer was stained with 0.5% crystal violet. (F) Viral plaque size. The diameters of viral plaques were measured using a scale in millimeters. Data indicate means ± SEM. 2-way ANOVA followed by multiple comparisons test, *** *p* < 0.001, **** *p* < 0.0001.

### SARS-CoV-2 D614G and Omicron BA.1 S present similar ability to inhibit interferon-beta (IFN-β) mediated luciferase activity

Given the reduced ability of the Omicron BA.1 strain to spread from cell-to-cell, we investigated the potential role of interferon (IFN) pathway in inhibiting viral replication. The SARS-CoV-2 S protein has been shown to inhibit the IFN pathway by antagonizing viral RNA pattern recognition receptor RIG-I signaling [25]. Using luciferase reporter assays we assessed the effect of D614G- and Omicron BA.1 S proteins on activation of the IFN-β and NF-κB signaling pathways. For this, HEK293T cells were transfected with plasmids encoding D614G- or Omicron BA.1 S proteins, or an empty vector, together with IFN-β or NF-κB Firefly luciferase reporter plasmids. At 24 hours post-transfection cells were stimulated with Sendai virus (SeV [Cantell strain], potent IFN inducer) or TNF-α (potent NF-κB pathway inducer) and incubated for 12 hours. After incubation Firefly luciferase activity was measured and used to assess and compare the inhibitory activity of the D614G- and Omicron BA.1-S proteins on these critical signaling pathways. Both D614G and Omicron BA.1 S proteins significantly (*p* < 0.01) inhibited SeV-induced IFN-β signaling, and no differences between the S variants was noted (Fig 3A). No inhibitory activity in the NF-κB signaling was observed (Figs 3B and 3C). Expression of D614G- and Omicron BA.1 S in transfected cells was confirmed by Western blot (Fig 3D). Importantly, results from the Western blot also demonstrated a lower cleavability of the precursor Omicron BA.1 S protein, as evidenced by markedly lower levels of the S2 subunit in Omicron BA.1 S transfected cell when compared to D614G S (Fig 3D). These findings suggest that the lower fusogenicity of the Omicron BA.1 S and the impaired ability of the rWA1-Omi-S virus to spread from cell-to-cell are likely due to a less efficient cleavage of S and further rule out the involvement of the innate IFN response on this important biological property.

**FIG 3.**
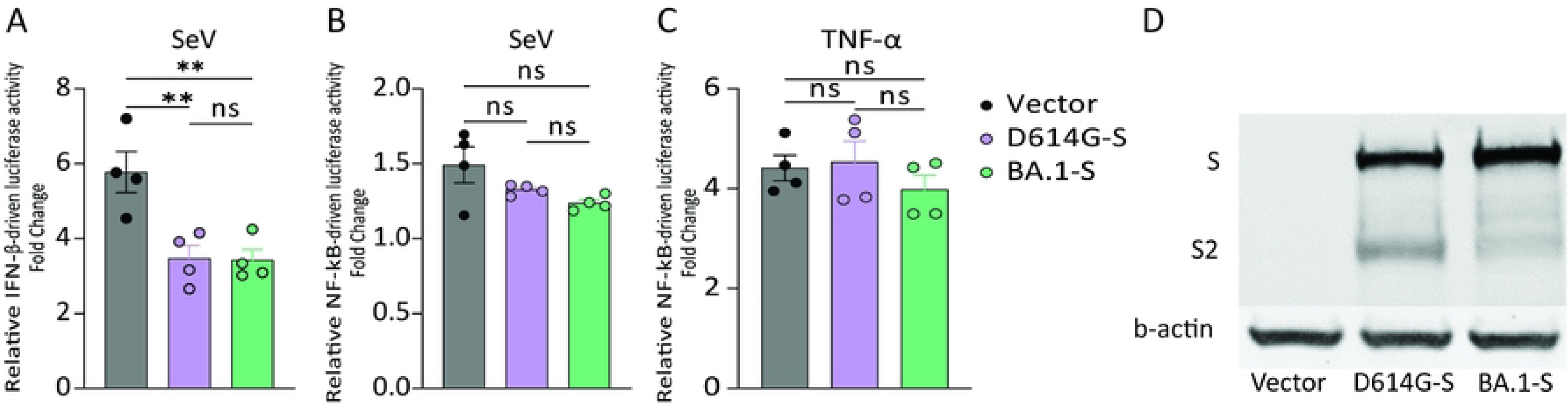
Innate immune inhibition by SARS-CoV-2 S protein. HEK293T cells were transiently transfected with plasmids encoding D614G and Omicron BA.1 S proteins together with IFN-β (A) or NF-κB (B-C) Firefly luciferase reporter plasmids. At 24 hours post-transfection, cells were then stimulated with SeV (A and B) or TNF-α (C) for 12 hours. Cell lysates were harvested, and the Firefly luciferase activity was measured using a luminometer. The ratio of luminescence obtained from the target reporters (IFN-β-Luc or NF-κB-Luc) to luminescence from the control Renilla reporter (pRN-Luc) was calculated to normalize the transfection efficiency. The relative IFN-β- or NF-κB-driven luciferase activity was calculated as fold change over the unstimulated cells. (D) HEK293T cells were transfected with plasmids encoding D614G and Omicron BA.1 S proteins and protein expression was detected by Western blot using an antibody against the S protein. B-actin was included as a loading control. (A-C) Data indicate mean ± SEM from four independent experiments. One-way ANOVA with multiple comparison test, ** *p* < 0.01, ns = not significant.

### The rWA1-Omi-S leads to subclinical infection and limited viral shedding in cats

The dynamics of infection, virus replication, and virus shedding of rWA1-D614G and rWA1-Omi-S were next assessed in cats. Following inoculation, clinical parameters, including rectal temperature, body weight, and clinical signs of respiratory disease, were monitored daily (Fig 4A). Animals inoculated with the rWA1-D614G became lethargic and presented increased body temperatures on days 2 and 3 post-infection (pi) (Fig 4B), whereas rWA1-Omi-S-inoculated cats remained subclinical. Additionally, rWA1-D614G-inoculated cats lost weight between days 1-7 pi, while all rWA1-Omi-S-inoculated cats gained weight throughout the 14-day experimental period (1 - 12% of their initial body weight) similar to the control mock-inoculated animals (Fig 4C).

**FIG 4.**
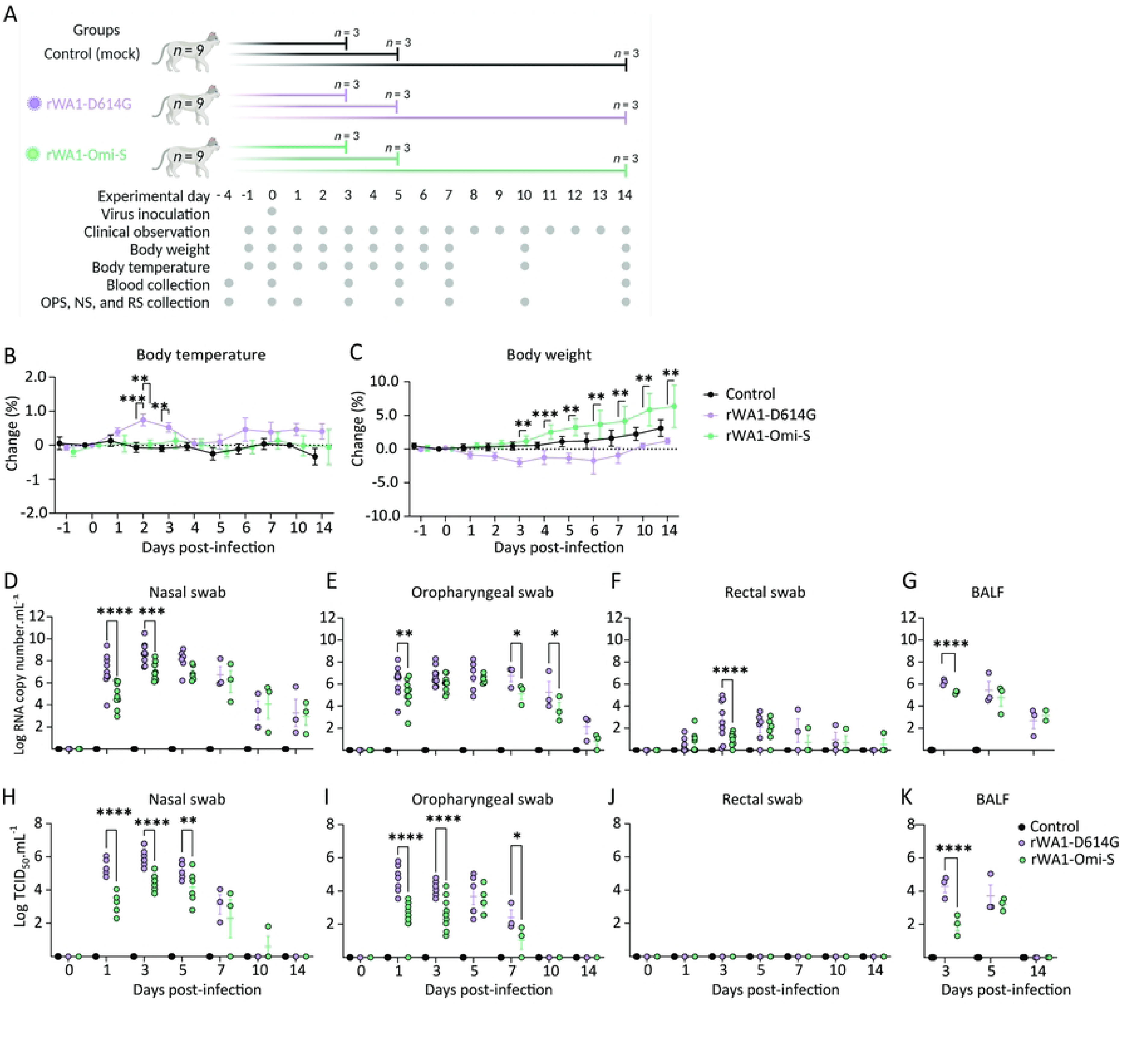
rWA1-Omi-S leads to subclinical infection and limited viral shedding in cats. (A) Experimental study design. Body temperature (B) and body weight (C) following intranasal inoculation of rWA1-D614G and rWA1-Omi-S throughout the 14-day experimental period. SARS-CoV-2 RNA loads quantified by rRT-PCR in nasal (D) and oral (E) secretions, and feces (F) collected on days 0, 1, 3, 5, 7, 10 and 14 pi and in bronchoalveolar lavage fluid (BALF) (G) collected on days 3, 5 and 14 pi. Infectious SARS-CoV-2 loads in nasal (H), oral (I) secretions, feces (J), and BALF (K) determined by virus titrations in rRT-PCR positive samples. Virus titers were determined using endpoint dilutions and expressed as TCID_50_.mL^-1^. The limit of detection (LOD) for infectious virus titration was 10^1.05^ TCID_50_.mL^-1^. Data indicate mean ± SEM of 3-9 animals per group per time point. 2-way ANOVA with multiple comparison test, * *p <* 0.05; ** *p <* 0.01; *** *p <* 0.005; **** *p <* 0.0001.

To assess virus replication and shedding dynamics of the recombinant SARS-CoV-2 following inoculation, nasal and oropharyngeal secretions and feces were collected using nasal (NS), oropharyngeal (OPS), and rectal swabs (RS). The samples were initially tested for the presence of SARS-CoV-2 RNA by real-time reverse transcriptase PCR (rRT-PCR). Viral RNA was detected between days 1 and 14 pi in NS and OPS secretions in the inoculated cats, with higher viral RNA loads detected between days 3 and 5 pi, which decreased thereafter through day 14 pi (Figs 4D and 4E). Viral RNA load was higher in nasal secretions of rWA1-D614G-inoculated animals when compared to rWA1-Omi-S-inoculated cats on days 1 and 3 pi (*p* < 0.001). Similarly, viral RNA load was higher in OPS secretions of rWA1-D614G-inoculated animals when compared to rWA1-Omi-S-inoculated cats on days 1, 7, and 10 pi (*p* < 0.05) (Fig 4E). Shedding of viral RNA in feces was markedly lower than in respiratory and oropharyngeal secretions and was characterized by intermittent detection of low amounts of viral RNA in feces, with rWA1-Omi-S-inoculated animals presenting lower RNA levels in feces on day 3 pi (*p* < 0.001) (Fig 4F). All control cats (mock-inoculated) remained negative throughout the 14-day experimental period.

The dynamics of infectious virus shedding were also assessed in NS, OPS, and RS for the presence of infectious virus. High viral loads were detected in NS and OPS secretions of rWA1-D614G-inoculated cats (Figs 4H and 4I). All rWA1-D614G-inoculated cats shed infectious SARS-CoV-2 between days 1–7 pi in nasal secretions, with viral titers ranging from 2.0 to 6.8 log TCID_50_.mL^-1^, whereas rWA1-Omi-S-inoculated animals shed significantly lower viral titers ranging from 1.8 to 5.5 log TCID_50_.mL^-1^ (Fig 4H). Cats inoculated with rWA1-D614G shed infectious virus between days 1–7 pi in the oropharyngeal secretions, with viral titers ranging from 1.8 to 5.8 log TCID_50_.mL^-1^, while rWA1-Omi-S-inoculated cats shed lower viral titers ranging from 1.3 to 4.55 log TCID_50_.mL^-1^ (Fig 4I). No infectious virus shedding was detected in feces in any of the groups (Fig 4J).

We also assessed the viral RNA load and infectious virus titers in bronchoalveolar lavage fluid (BALF). Three cats from each group (control, rWA1-D614G, rWA1-Omi-S) were humanely euthanized on days 3, 5, and 14 pi (Fig 4A). Viral RNA was detected in BALF of all inoculated animals on days 3 and 5 pi (Fig 4G). All control cats (mock-inoculated) tested negative by rRT-PCR. On day 3 pi, infectious virus titers in BALF of cats infected with rWA1-D614G varied from 3.5 to 5.0 log TCID_50_.mL^-1^, whereas rWA1-Omi-S-inoculated animals, presented lower infectious virus titers ranging from 1.3 and 2.5 log TCID_50_.mL^-1^ (*p* < 0.001) (Fig 4K). On day 5 pi, viral titers in BALF from cats infected with rWA1-D614G varied from 3.0 to 5.0 log TCID_50_.mL^-1^, while rWA1-Omi-S-inoculated cats were 2.8 to 3.5 log TCID_50_.mL^-1^ (Fig 4K). Viral RNA was detected in BALF from all inoculated cats on day 14 pi, (Fig 4G), however, no infectious virus was detected (Fig 4K). Together, these results demonstrate that the rWA1-Omi-S presents lower pathogenicity and replication ability when compared to the rWA1-D614G recombinant virus in cats.

### rWA1-Omi-S presents reduced replication in tissues

The tissue tropism and replication sites of rWA1-D614G and rWA1-Omi-S were assessed following intranasal inoculation in cats. For this, nasal turbinate, palate/tonsil, retropharyngeal lymph node, trachea, lung, mediastinal lymph node (LN), heart, liver, spleen, kidney, small intestine, and mesenteric LN were collected from three cats per group on days 3, 5 and 14 pi following euthanasia and processed for rRT-PCR, infectious virus titration, and *in situ* hybridization and immunofluorescence staining. SARS-CoV-2 RNA was detected in several tissues sampled from each group, with higher viral RNA loads being detected on days 3 and 5 pi (Figs 5A-5C). The highest viral RNA loads were detected in the nasal turbinate on day 3 and 5 pi in rWA1-D614G-inoculated animals, with lower viral RNA loads detected in tissues from rWA1-Omi-S-inoculated cats (Figs 5A and 5B). To assess virus replication in tissues, subgenomic viral RNA (sgRNA) was quantified by qRT-PCR targeting the envelop (E) gene. sgRNA was consistently detected in nasal turbinate, palate/tonsil, trachea, retropharyngeal LN, and lung from rWA1-D614G-inoculated cats on days 3 and 5 pi (Figs 5D and 5E). The highest sgRNA loads were observed in nasal turbinate on days 3 and 5 pi (Figs 5D and 5E). sgRNA was detected in nasal turbinate, soft palate/tonsil, retropharyngeal LN, trachea, and lung from rWA1-D614G-inoculated, whereas sgRNA was detected only in nasal turbinate and trachea from rWA1-Omi-S-inoculated cats on day 3 pi (Fig 5D). On day 5 pi, sgRNA was detected in nasal turbinate, soft palate/tonsil, retropharyngeal LN, trachea, lung, and in mesenteric LN from 1 out of 3 cats regardless of the virus inoculated (Fig 5E). On day 14 pi, viral RNA loads decreased when compared to early time points and sgRNA load was higher in retropharyngeal LN of rWA1-D614G-inoculated cats when compared to rWA1-Omi-S-inoculated animals (*p* < 0.001) (Fig 5F). All tissues from the control animals (mock-inoculated) tested negative for viral RNA by RT-PCR (Figs 5A-5F).

**FIG 5.**
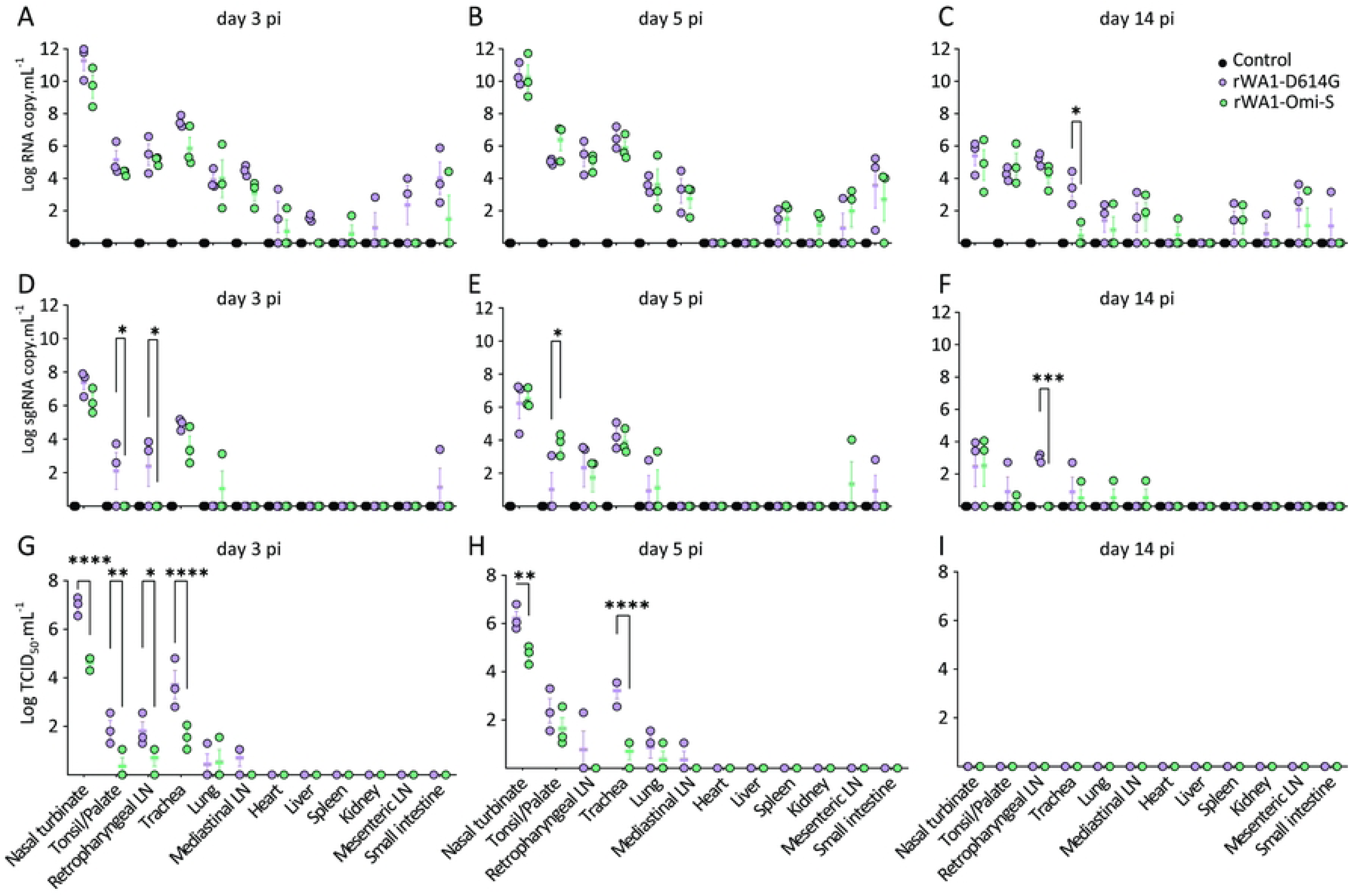
rWA1-Omi-S presents reduced replication in tissues. SARS-CoV-2 RNA loads assessed by rRT-PCR in tissues of cat collected on days 3 (A), 5 (B) and 14 (C) pi. Subgenomic SARS-CoV-2 RNA (sgRNA) loads (replicating RNA) assessed by rRT-PCR in tissues of cats collected on days 3 (D), 5 (E), and 14 (F) pi. Infectious SARS-CoV-2 in tissues assessed by virus titration in rRT-PCR positive tissue samples collected on days 3 (G), 5 (H), and 14 (I) pi. Virus titers were determined using endpoint dilutions and expressed as TCID_50_.mL^-1^. The limit of detection (LOD) for infectious virus titration was 10^1.05^ TCID_50_.mL^-1^. Data indicate mean ± SEM of three animals per group per time point. 2-way ANOVA with multiple comparison test, * *p <* 0.05; ** *p <* 0.01; *** *p <* 0.005; **** *p <* 0.0001.

The presence and titers of infectious SARS-CoV-2 in tissues were next assessed by virus titrations in rRT-PCR positive tissues. Detection of infectious virus and infectious viral loads were consistent with detection of sgRNA (Figs 5D-5I). Infectious SARS-CoV-2 was consistently detected in respiratory tract tissues on days 3 and 5 pi including in nasal turbinate, palate/tonsil, retropharyngeal LN, trachea, and lung from cats from both groups (Figs. 5G and 5H). Viral loads, however, were higher in tissues from rWA1-D614G-inoculated cats. The highest viral titers were observed in the nasal turbinate (titers ranging 5.8 to 7.3 log TCID_50_.mL^-1^) from rWA1-D614G-inoculated cats on days 3 and 5 pi, while markedly lower viral loads (4.0 to 5.0 log TCID_50_.mL^-1^) were detected in rWA1-Omi-S inoculated cats (Figs 5G and 5H). Interestingly, although on day 5 pi infectious virus titers detected in rWA1-D614G-inoculated animals were slightly lower than in day 3 pi, viral titers in tissues from rWA1-Omi-S-inoculated cats were slightly higher than viral titers detected on day 3 pi (Figs 5G and 5H). Together these results indicate reduced replication of the rWA1-Omi-S in the respiratory tract and associated lymphoid tissues when compared to the rWA1-D614G recombinant virus in cats.

### rWA1-Omi-S presents reduced replication and induces milder pathological changes in the respiratory tract of cats

The tissue distribution of SARS-CoV-2 in the respiratory tract and associated lymphoid tissues was assessed by *in situ* hybridization (ISH) and IFA staining. For this, nasal turbinate and lung from all animals were examined by ISH using the RNAscope^®^ ZZ technology and by IFA using a SARS-CoV-2 N specific antibody. While intense labeling for viral RNA and staining for the viral N protein were consistently observed in nasal turbinate from rWA1-D614G-inoculated cats, only modest and sporadic staining was detected in tissues from rWA1-Omi-S-inoculated animals on days 3 and 5 pi (Figs 6A and 6B). When we compared the intensity and distribution of both viral RNA and N protein in nasal turbinate from the two inoculated groups, the lower infectivity of the rWA1-Omi-S virus was evident by markedly lower levels of viral RNA and N detection across all animals tested (Figs 6A and 6B).

**FIG 6.**
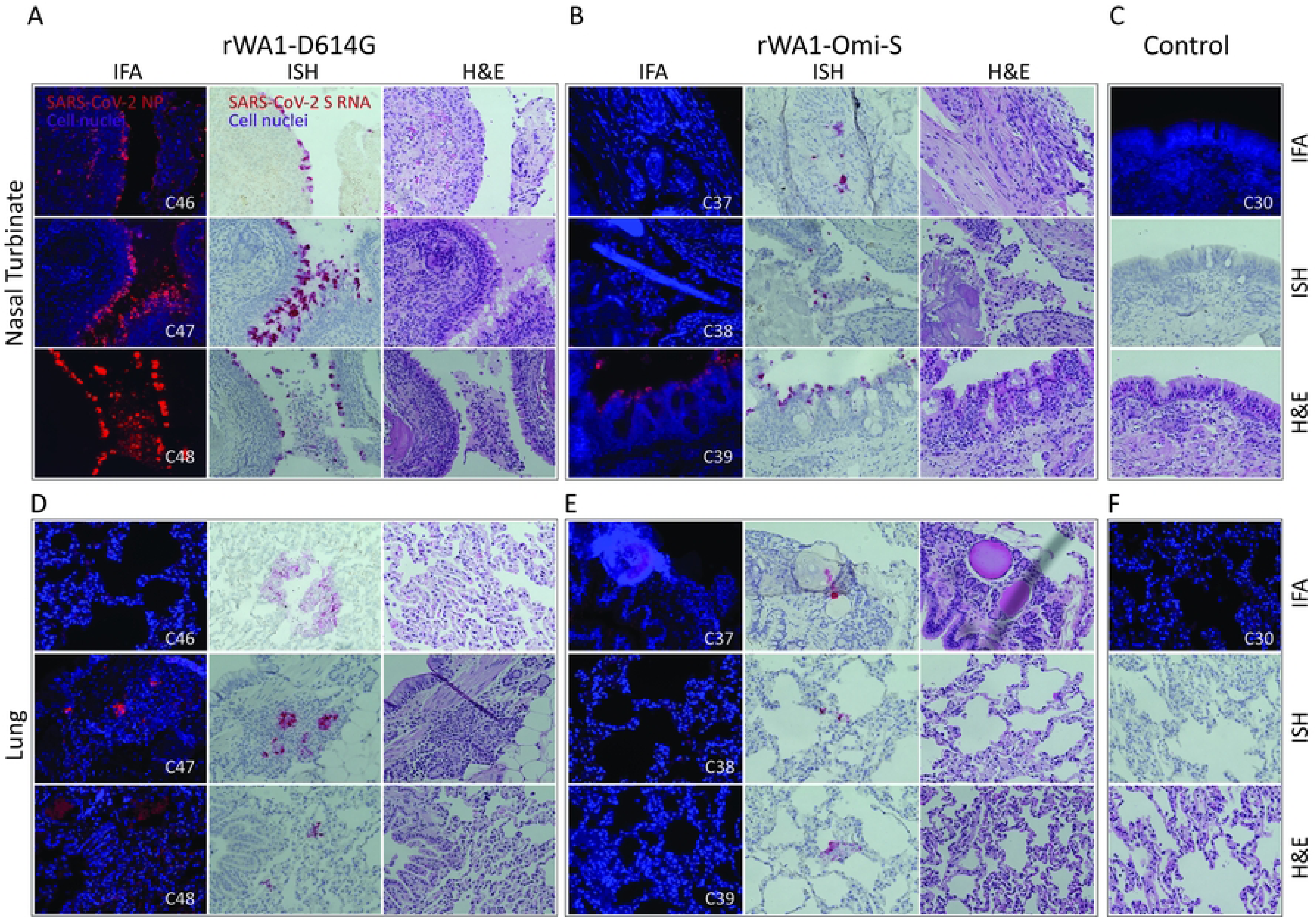
rWA1-Omi-S presents reduced replication in the respiratory tract of cats. Immunofluorescence (IFA), *in situ* hybridization (ISH), and hematoxylin and eosin (HE) staining in nasal turbinate (A-C) and lung (D-F) of cats inoculated with rWA1-D614G and rWA1-Omi-S, and control cats. Nasal turbinate and lungs were collected from rWA1-D614G (A and D) and rWA-1-Omi-S (B and E) or mock (C and F) inoculated cats on day 3 pi. The IFA was performed using a monoclonal antibody against N protein of SARS-CoV-2 (red labelling). In ISH, the viral RNA (red labelling) was performed using a probe targeting the SARS-CoV-2 S RNA. Note, both IFA and ISH showed intense labelling on tissues from rWA1-D614G-inoculated cats and less abundant labelling on tissues from rWA1-Omi-S-inoculated cats. Representative images of nasal turbinate and lungs collected from three inoculated cats with rWA1-D614G (A and D) or rWA1-Omi-S (B and E) and one mock-inoculated cat are shown.

Epithelial cells of the nasal mucosa were the predominant cell type positive for SARS-CoV-2 in the rWA1-D614G-inoculated animals, with extensive virus labeling observed on day 3 pi. Additionally, middle and basilar areas of the epithelium were also sporadically stained (Fig 6A). In the lung, sparse staining of bronchial cells was observed on days 3 and 5 pi (Fig 6D). Staining in the lung was more frequent in the interstitial regions especially in cells of the bronchiolar glands. In rWA1-Omi-S-inoculated cats viral RNA and N protein staining in both the nasal turbinate and lung, were consistently lower (number of cells and extension of staining) than that observed in tissues from rWA1-D614G-inoculated animals (Figs 6A, 6B, 6D, and 6E). No viral RNA or N protein staining was observed in tissues from control animals (Figs 6C and F). These results demonstrate that the rWA1-Omi-S virus presents limited tissue distribution, replication and spread in the respiratory tract when compared to rWA1-D614G in cats.

Histological examination was performed on tissues collected on days 3, 5 and 14 pi. The rWA1-D614G-inoculated cats presented with aggregates of fibrin, cellular debris and leukocytes partially filling the nasal passages on days 3 (Fig 6A) and 5 pi. Marked dislodging of nasal epithelial cells and mild to moderate epithelial necrosis were also noticed. In the lungs, bronchiolar necrosis with accumulation of exudates in the lumen containing necrotic epithelial cells and mononuclear inflammatory cells was observed on days 3 and 5 pi. Mononuclear infiltration was also noticed in the sub-mucosa of bronchi and bronchiole on days 3 (Fig 6D) and 5 pi. Thickening of the alveolar septa was consistently noticed at all time points examined with abundant mononuclear cellular infiltration on day 3 pi (Fig 6D).

In general, histological changes observed in rWA1-Omi-S-inoculated cats were milder than those observed in rWA1-D614G-infeted animals. Deciliation of the nasal epithelial cells and infiltration of mononuclear cells were observed in the submucosa of the nasal passage from rWA1-Omi-S inoculated cats (Fig 6B). Small accumulation of fibrin and cellular aggregates in the nasal passage was also noticed on days 3 and 5 pi. In the lungs, mild exudation was noticed in bronchiolar lumen on days 3 and 5 pi. Thickening of alveolar septa with mononuclear infiltration was noticed on day 3 (Fig 6D) and 5 pi. All control cats presented normal histology in the nasal passages (Fig 6C) and lungs (Fig 6F). Together, these results demonstrate that the rWA1-Omi-S presents a limited pathogenicity in the infected cats compared to rWA1-D614G.

### Antibody responses following rWA1-D614G and rWA1-Omi-S infection in cats

The antibody responses to SARS-CoV-2 were assessed by luciferase immunoprecipitation (LIPS) and virus neutralization (VN) assays. We used the LIPS assay to determine the kinetics of antibody responses against the N protein in the sera of the rWA1-D614G- and rWA1-Omi-S-infected cats. Serum samples from control and rWA1-D614G- and rWA1-Omi-S-infected-inoculated cats were collected on days 0, 7, and 14 pi and tested by LIPS assay (*n* = 3 cats per group) [26]. None of the control or inoculated cats reacted strongly to the NLuc-tagged N protein on days at day 0 and 7 pi (Fig 7A). At 14 dpi, serum from both rWA1-D614G- and rWA1-Omi-S-inoculated animals showed strong binding to the NLuc-tagged N protein antigen confirming presence of N protein specific antibodies, with rWA1-D614G-infected cats presenting higher antibody levels as evidenced by higher luminescence than the rWA1-Omi-S-inoculated cats (Fig 7A).

**FIG 7.**
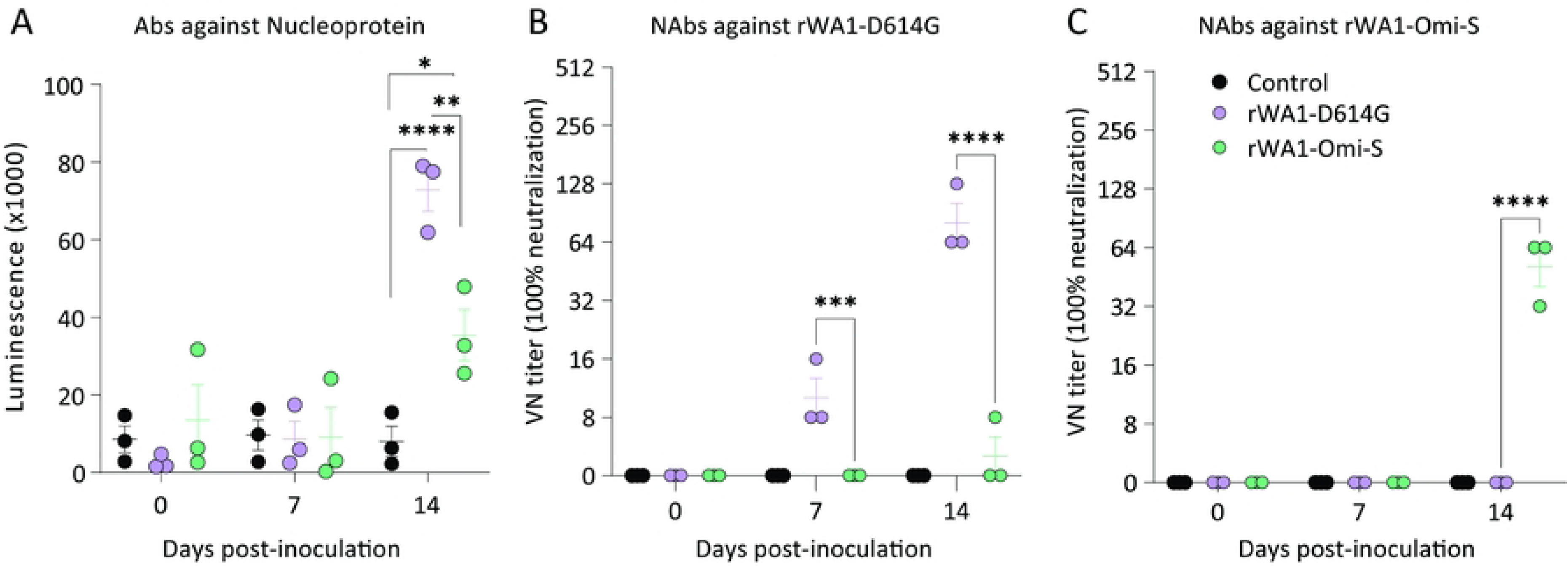
Antibody responses following rWA1-D614G and rWA1-Omi-S infection in cats. The kinetics of antibody responses against N protein measured by LIPS assay (A). NAb responses to SARS-CoV-2 and cross-reactivity between rWA1-D614G and rWA1-Omi-S recombinants were assessed by VN (100% neutralization) (B and C). Data indicate mean ± SEM of three animals per group per time point. 2-way ANOVA with multiple comparison test, * *p <* 0.05; ** *p <* 0.01; *** *p <* 0.005; **** *p <* 0.0001.

Neutralizing antibodies (NAbs) responses to SARS-CoV-2 and cross-reactivity between rWA1-D614G and rWA1-Omi-S were evaluated. All rWA1-D614G-inoculated cats presented NAbs titers as early as day 7 pi, while NAbs in rWA1-Omi-S-inoculated cats were only detected on day 14 pi. Neutralizing responses to the homologous virus were significantly higher than the cross-neutralizing responses (*p* < 0.001) (Figs 7B and 7C). NAbs titers in rWA1-D614G serum ranged from 64 to 128 against rWA1-D614G virus, whereas no NAbs titers detected against rWA1-Omi-S (titer lower than 8). rWA1-Omi-S infection induced lower and delayed NAbs, which were efficient in neutralizing SARS-CoV-2 (Fig 7C). All control animals remained seronegative throughout the experiment. These results confirm seroconversion of all rWA1-D614G- and rWA1-Omi-S-inoculated cats, demonstrating that all the inoculated animals were infected. Moreover, results showed limited cross-neutralizing activity between rWA1-D614G and rWA1-Omi-S serum.

## Discussion

Here, we demonstrated that the S protein plays an important role in SARS-CoV-2 virulence and is a major determinant of the attenuated phenotype of SARS-CoV-2 Omicron BA.1 variant. In cell culture systems, the Omicron BA.1 S seems to mediate efficient entry into host cells, however, the rWA1-Omi-S presented reduced fusogenicity and ability to spread from cell-to-cell when compared to the B.1 D614G S bearing rWA1-D614G. In cats, the rWA1-Omi-S presented subclinical infection, reduced replication in the upper respiratory tract, and decreased virus shedding and pathology, when compared to the ancestral rWA1-D614G virus, indicating that the S protein is as a major determinant of SARS-CoV-2 pathogenicity.

The S protein of the Omicron BA.1 variant possesses 36 mutations compared to the ancestral D614G lineage. Several of the affected residues modulate receptor binding, protease cleavage efficiency, and cell-to-cell fusion, and contribute to virus replication and escape from NAbs. The S protein binds to the hACE2 receptor and mediates entry into susceptible host cells. Studies showed that the S protein of Omicron variant utilizes the hACE2 receptor with enhanced efficiency than ancestral SARS-CoV-2 strains, a feature that has been speculated to contribute to rapid transmission of the virus [18,27–29]. Omicron BA.1 S protein presents an enhanced RBD-ACE2 binding interface through the interactions of amino acids N501Y, Q498R, and T478K with the host receptor in target cells. Notably, these mutations are reminiscing features of other variants, including Alpha (N501Y) and Delta (L452R and T478K) [29]. Additional mutations such as Q493R and S477N also contribute to enhanced S-ACE2 interactions [18,30]. Notably, results of the infectivity assays in the present study using infectious recombinant viruses (rWA1-D614G and rWA1-Omi-S) demonstrated a more efficient entry of the Omicron BA.1 S harboring virus when compared to the ancestral D614G S virus. As the infection progressed, however, the rWA1-D614G virus, owing to its enhanced fusogenic activity and ability to spread from cell-to-cell, overtook the rWA1-Omi-S-infection resulting in higher spread and replication efficiency of the virus.

Recent studies demonstrated that the Omicron variant BA.1 is less dependent on transmembrane serine protease 2 (TMPRSS2), less efficient in S cleavage, less fusogenic, and favors the endosomal pathway for virus entry [20,28,31]. In line with this, results here with the Omicron S protein and the infectivity assays with the rWA1-Omi-S also showed reduced fusogenicity and syncytia formation in hACE2_BHK21 and cACE2_BHK21 cells as compared to the D614G S and the rWA1-D614G, indicating that S protein of Omicron contributes to this phenotype. The lower fusogenicity of the Omicron S could also affect the ability of the virus to spread from cell-to-cell. This was confirmed by the smaller plaque size observed for the rWA1-Omi-S compared to the rWA1-D614G. S mutations H655Y and T547K have been shown to contribute to the low fusogenicity of Omicron variants, while the H655Y mutation also dictates the enhanced endosome entry pathway utilization of Omicron [20,32]. SARS-CoV-2 S antagonizes viral RNA recognition by RIG-I blocking the induction of IFN and downstream IFN stimulated genes (ISGs) [25]. Subsequent studies showed that the S protein interacts with signal transducer and activator of transcription 1 (STAT1) to block its association with Janus kinase 1 (JAK1) [33]. Considering about 36 amino acid mutations in the S protein between Omicron BA.1 and B.1 D614G variants, we examined if these S proteins modulate innate immune inhibitory pathways differently. Differential modulation of innate responses and more specifically of IFN pathways could, for example, account for the reduced plaque size and lower ability of the rWA1-Omi-S virus to spread from cell-to-cell. Results from an IFNβ-driven luciferase reporter assay, however, revealed comparable inhibition of the IFN-β reporter activity by B.1 D614G and Omicron BA.1 S proteins. Together these results rule out impaired innate immune modulation as a factor contributing for the lower replication ability and cell-to-cell spread observed for rWA1-Omi-S virus. These observations further point towards the decreased cleavability and fusogenic activity of the Omicron BA.1 S as a major driving force contributing to the lower overall infectivity and ability of the virus to spread from cell-to-cell.

The Omicron BA.1 variant is more immune evasive and less virulent than the other major identified variants [10,12–15,18,23,27,28,30,31,34–36]. In Syrian hamsters, the SARS-CoV-2 B.1 D614G produced significant illness, higher viral loads in secretions and tissues, while the Omicron BA.1-inoculated animals were subclinical and had reduced virus replication and shedding [37]. Other studies in hamsters and humanized K18 transgenic hACE2 mice also showed lower levels of lung infection and replication, decreased clinical disease and pathology in Omicron inoculated animals when compared to historical isolates or other SARS-CoV-2 variants [23,31,34]. Similarly, using domestic cats, a naturally susceptible animal species, we recently showed that Omicron BA.1.1 causes lower pathogenicity than ancestral B.1 and Delta variants [24]. Here by using recombinant chimeric viruses (rWA1-D614G and rWA1-Omi-S) we sought to determine and dissect the role and contribution of the S protein for the attenuated disease phenotype of Omicron BA.1. For this, we inoculated cats with rSARS-CoV-2 carrying Omicron S (rWA1-Omi-S) and D614G S (rWA1-D614G) in the backbone of ancestral WA1 isolate. While rWA1-D614G inoculated cats were febrile, lethargic, and lost or maintained steady body weight, rWA1-Omi-S inoculated cats remained subclinical and gained weight throughout the experiment. Similarly, the rWA1-Omi-S presented significantly lower replication in the respiratory tract and associated lymphoid tissues of inoculated animals, as evidenced by lower virus shedding and milder viral replication and pathological changes in target respiratory organs. These results confirm a major role for the Omicron BA.1 S for the attenuation of the virus in the feline model of infection. However, the role and contribution of Omicron-specific mutations that fall outside of S protein for virus infection and disease pathogenesis cannot be formally ruled out. In fact, recent studies using K18 transgenic hACE2 mice showed that S contributes to the attenuated phenotype of Omicron BA.1 and mutation outside S protein are also likely to contribute to the virulence of the virus in this mouse model [17,38]. In these studies, intranasal inoculation of K18 transgenic hACE2 mice with Omicron BA.1 caused no significant body weight loss, while the rWA1 bearing Omicron S caused less severe disease than the ancestral virus but was not as attenuated as the natural Omicron [17]. Survival assessments revealed non-fatal infection in Omicron-infected mice compared to the 100% and 80% mortality in ancestral D614G and Omi-S infected mice [17]. Another study in K18 transgenic hACE2 mice also showed an S independent attenuation of Omicron variant [38]. While the ancestral WA1 virus bearing the Omicron S protein was lethal, the Omicron virus bearing WA1 S protein was attenuated in K18 transgenic hACE2 mice [38]. More recently, another study showed that while Omicron BA.1 and BA.2 cause minimal to negligible morbidity and mortality in K18 hACE2 mice, animals infected with Omicron BA.5 exhibited high weight loss and mortality that was dependent on the age of the mice [39].

Mutations in the S protein of the Omicron variants, particularly in the RBD, lead to widespread escape from NAb responses causing vaccine breakthrough infection and increase transmission [10,11,13–15,28]. As a result, Omicron showed lower neutralizing sensitivity to antibodies elicited by earlier variants or by immunization [10,11,15,28]. NAbs were detected in serum of rWA1-D614G infected cats as early as day 7 pi, while were only detected on day 14 pi for the rWA1-Omi-S. Cross-neutralization assay showed no cross-neutralizing activity between rWA1-D614G and rWA1-Omi-S serum, confirming that the S protein contributes to the neutralizing activity and the Omicron S extensively escapes neutralization by antibodies elicited against ancestral WA1-D614G S. Our previous study also showed no or little cross-reactivity in serum neutralization between SARS-CoV-2 B.1 and Omicron BA.1.1-infected cat serum [24].

In summary, our study demonstrates the critical role and contribution of the Omicron BA.1 S protein for virus infectivity, cell-to-cell spread and replication *in vitro* and *in vivo*. Our *in vitro* studies show that Omicron BA.1 S presents a lower fusogenicity ability when compared to the ancestral D614G S, which leads to decreased cell-to-cell spread of the virus. These results were confirmed *in vivo* in a feline model of infection. Importantly, while other non-S mutations may also contribute to the lower pathogenicity of Omicron BA.1 variant, our results demonstrate that the S is a major player contributing to virulence and pathogenesis of SARS-CoV-2.

## Methods

### Cells and viruses

Vero E6 (ATCC^®^ CRL-1586™), and Vero E6/TMPRSS2 (JCRB Cell Bank, JCRB1819) were cultured in Dulbecco’s modified eagle medium (DMEM), supplemented with 10% fetal bovine serum (FBS), L-glutamine (2mM), penicillin (100 U.mL^−1^), streptomycin (100 μg.mL^−1^) and gentamicin (50 μg.mL^−1^). BHK21 and HEK293T cells were maintained in complete growth media consisting of Minimum Essential Media (Corning, 10-010-CV), supplemented with 10% FBS, and supplemented with penicillin (100 U.mL^−1^) and streptomycin (100 μg.mL^−1^). The cell cultures were maintained at 37°C with 5% CO_2_. Sendai virus (SeV) (Cantell strain) was propagated in 11-days-old embryonated chicken eggs and titrated using hemagglutination assay (HA).

The bacterial artificial chromosome (BAC) harboring the whole genome of the ancestral SARS-CoV-2 WA1-D614G (rWA1-D614G) and S gene of Omicron BA.1 variant in the backbone of the ancestral rWA1-D614G (rWA1-Omi-S) were generated as previously described [40]. Recombinant viruses were rescued under BSL-3 laboratory conditions using Vero E6 TMPRSS2. Briefly, Vero E6 TMPRSS2 cells were seeded in 6-well plates (3×10^5^ per well) and transfected with 2 µg/well of the respective SARS-CoV-2 BAC DNA (pBAC-WA1-D614G or pBAC-WA1-Omi-S [BA.1 S]) using Lipofectamine 3000. After 24 hours, culture medium was replaced with complete growth media containing 5% FBS and incubated for an additional 48 hours. The culture supernatant was collected and labelled as P0. The recombinant viruses rWA1-D614G and rWA1-Omi-S were passaged twice in Vero E6 TMPRSS2 cells. The whole genome sequences of the virus stocks were determined to confirm that no mutations occurred during rescue and amplification in cell culture and the sequences were deposited in the GenBank (accession number: OR626632-OR626633). The titers of virus stocks were determined by plaque assays, calculated according to the Spearman and Karber method, and expressed as plaque-forming units per milliliter (PFU.mL^−1^). Sequenced verified viral stocks were used in all experiments described in this study.

### Generation of ACE2 stably expressing cell lines

Human and cat ACE2 expressing lentiviral plasmids (pscALPS-hACE2 and pscALPS-cACE2) were obtained from Addgene (158081 and 158082, respectively). Each gene was tagged with a c-terminal Myc-tag epitope to facilitate detection. Lentiviral particles encoding hACE2 and cACE2 were generated in HEK293T cells. For this, 7.2 × 10^5^ cells were seeded in each well of a 6 well plate and allowed to adhere overnight. Each well was then transfected with the lentiviral packaging vectors psPAX2 (1 µg, Addgene, 12260), and pMD2.G (1 µg, Addgene, 12259), plus the pscALPS-hACE2 or -cACE2 (1 µg) or the empty pscALPS_puro (Addgene, 128504) vector as a negative control. Lentiviral particles were harvested from the supernatant of transfected cells at 72 hours post-transfection, cleared by centrifugation, aliquoted and stored at −80°C until use.

Target BHK21 cells in exponential growth phase were seeded 7.2 × 10^5^ cells in a 6 well plate and allowed to adhere overnight. Prior to transduction, aliquots of the lentivirus were brought to room temperature and gently inverted to mix. Media was removed from the BHK21 cells and 1 mL of each lentivirus was added per well and adsorbed for 2 hours at 37 °C with plate rocking every 30 minutes. After adsorption, 2 mL of complete growth media was added, and cells were incubated for 72 hours. At 72 hours media was replaced with complete growth media containing 5 µg.mL^-1^ of Puromycin Dihydrochloride for selection (Gibco, A1113803). Following complete death of non-transduced cells, expression of hACE and cACE2 was validated via immunofluorescence and immunoblots through detection of the c-terminal myc tag with a 1:500 dilution of antibody Myc-Tag-9B11 Mouse mAb (Cell Signaling Technology, 2276).

### Ethics statement

The protocols and procedures for generation of the recombinant viruses were reviewed and approved by Cornell University Institutional Biosafety Committee (MUA 16373-2). All animals were handled in accordance with the Animal Welfare Act. The study procedures were reviewed and approved by the Institutional Animal Care and Use Committee at Cornell University (IACUC approval number 2020-0064).

### SARS-CoV-2 infectivity assay

To assess infectivity of rWA1-D614G and rWA1-Omi-S viruses, hACE2_BHK21, cACE2_BHK21, Vero E6 and Vero E6 TMPRSS2 cells were seeded into 8-well glass chamber slides (Millipore, PEZGS0816). At 24 hours when cells were approximately 80% confluent, they were inoculated with rWA1-D614G and rWA1-Omi-S at a multiplicity of infection (MOI) of 1 and incubated at 4°C for 1 hour. Following incubation, the inoculum was removed, 1 mL of complete growth media was added to each well and plates were incubated at 37°C with 5% CO_2_. At 4-, 8-, 12- and 24-hours post-inoculation cells were fixed with 3.7% PFA for 30 minutes and washed with PBS. To monitor SARS-CoV-2 entry, infectivity and spread cells were stained with a SARS-CoV-2 anti-N rabbit polyclonal antibody for 1 hour followed by 594-conjugated secondary Goat anti-Rabbit IgG (VWR, GtxRb-003-D594NHSX) for 1 hour. Nuclei were stained with 4′,6-diamidino-2-phenylindole (DAPI) (ThermoFisher Scientific, 62248). The percentage of infected cells at each time point was quantified using ImageJ software.

### Fusogenicity Assay

To assess and compare the fusogenicity of D614G and Omi-S S proteins, hACE2_BHK21, cACE2_BHK21, or control BHK21 cells were seeded into 8-well glass chamber slides (Millipore, PEZGS0816) to a final confluency of 80%. Initially, cells were transfected with 250 ng of plasmids encoding the SARS-CoV-2 S genes delivered by pTwist-SARS-CoV-2 Δ18 D614G (Addgene, 164437) or pTwist-SARS-CoV-2 Δ18 B.1.1.529 (Addgene, 179907), incubated overnight at 37°C with 5% CO_2_ and fixed with 3.7% PFA for 20 minutes. To assess the S fusogenicity during infection hACE2_BHK21, cACE2_BHK21, or control BHK21 cells were seeded in 8-well glass chamber slides (Millipore, PEZGS0816) to a final confluency of 80% and inoculated with rWA1-D614G or rWA1-Omi-S viruses (MOI = 0.1). Viruses were adsorbed to cells for 1 hour at 4°C after which the inoculum was removed and 1 mL complete growth media added to the cells. Cells were incubated for 24 hours at 37°C with 5% CO_2_ and fixed with 3.7% PFA for 30 minutes. After fixation, transfected and infected cells were stained with an anti-N rabbit polyclonal antibody followed by incubation with secondary 594-conjugated Goat anti-Rabbit IgG (VWR, GtxRb-003-D594NHSX). Nuclei were stained with DAPI (ThermoFisher Scientific, 62248). For imaging, PBS was removed from all wells, the outer well walls were removed, and the glass slides were mounted with mounting media (Ibidi, 50-305-707) and coverslips. Representative images were taken across three replicates at 10X, and individual nuclei were counted using ImageJ. The fusogenicity of each S protein was quantitated for each of the three replicate images using the formula: percentage of nuclei incorporated into syncytia = (# of syncytia enclosed nuclei) / (total # of nuclei in field).

### Plaque size determination

The ability of rWA1-D614G and rWA1-Omi-S to spread from cell-to-cell were evaluated using plaque assays. For this, Vero E6, hACE2_BHK21, and cACE2_BHK21 cells were seeded in 6-well plates (3×10^5^ cells per well) and at 24 hours were infected with rWA1-D614G and rWA1-Omi-S (10 plaque forming units [PFU] per well). After 1 hour incubation at 37°C, the inoculum was removed and 2 mL of media containing 2X complete growth media and 1% SeaKem agarose (final conc. 1X media 0.5% agarose) was added to each well. Once agarose polymerized, the plate was transferred to the incubator 37°C 5% CO_2_ for 72 h. The agarose overlay was removed, cells were fixed with 3.7% formaldehyde solution for 30 minutes and stained with 0.5% crystal violet solution for 10 minutes at room temperature. The images and plaque size were determined using a Keyence BZ-X810 Microscope.

### Viral growth kinetics

The replication kinetics of rWA1-D614G and rWA1-Omi-S were examined *in* Vero E6 (120,000 cells per well), hACE2_BHK21, and cACE2_BHK21 (360,000 cells per well) cells seeded in 12-well plated and incubated at 37°C for 24 hours until they reached 80 - 90% confluency. Cells were inoculated with rWA1-D614G and rWA1-Omi-S with a MOI of 0.1 (500 µL per well) and incubated at 4°C for 1 hour. Following incubation, the inoculum was removed, and 1 mL of complete growth media was added per well and cells incubated at 37°C with 5% CO_2_. Cells and supernatant were harvested at 4, 8-, 12-, 24- and 48-hours post-inoculation and stored at −80°C. Time point 0 hour was an aliquot of virus inoculum stored −80°C as soon as inoculation was completed. Virus titers were determined in Vero E6 TMPRRSS2 cells on each time point using end-point dilutions and the Spearman and Karber’s method and expressed as TCID_50_.mL^−1^.

### Luciferase reporter assay

The ability of the D614G and Omi-S to modulate innate immune pathway activation was investigated using luciferase assays. For this, HEK293T cells were seeded in 6-well plate at 2×10^5^ cells per well and transfected with pIFN-β-Luc or pNF-κB-Luc (200 ng/well) and pRN-Luc (50 ng) reporter plasmids in combination with S protein expressing plasmids pTwist-SARS-CoV-2 Δ18 D614G (Addgene, 164437) or pTwist-SARS-CoV-2 Δ18 B.1.1.529 (Addgene, 179907) or a pTwist empty vector (250 ng) using Lipofectamine 3000 (Invitrogen, L3000001). At 24 hours post-transfection, cells were stimulated with SeV, Cantell strain (100 hemagglutination units per well) or TNF-α (25 ng/well) for 12 hours. After stimulation cells were lysed and luciferase activity measured using the dual luciferase assay (Promega, E2940). Luminescence was measured using a luminometer plate reader (BioTek Synergy LX Multimode Reader). The ratio of luminescence obtained from the target reporters (IFN-β-Luc or NF-κB-Luc) to luminescence from the control Renilla reporter (pRN-Luc) was calculated to normalize the transfection efficiency. Then, the relative IFN-β- or NF-κB-driven luciferase activity was calculated as fold change over the unstimulated cells. Confirmation of the S expression in HEK293T transfected cells was performed by Western blot using antibody against SARS-CoV-2 S2 protein (Sino Biological).

### Animals housing and experimental design

A total of twenty-seven 20-65-month-old domestic cats (*Felis catus*) (five females and four males [n = 9] per group) were obtained from Clinvet (Waverly, NY, USA). Animals were donated to Cornell University to support the reduction of animal use in research. All animals were housed in the animal biosafety level 3 (ABSL-3) facility at the East Campus Research Facility (ECRF) at Cornell University. After acclimation, and prior to virus inoculation, nasal swabs and serum were collected from all animals and tested by rRT-PCR for SARS-CoV-2 and serum neutralization using SARS-CoV-2 B.1 and Omicron lineages which confirmed that the animals were negative for SARS-CoV-2 [24]. On day 0, cats were anesthetized and inoculated intranasally with 1 mL (0.5 mL per nostril) of a virus suspension containing 5 × 10^5^ PFU of rWA1-D614G or rWA1-Omi-S. Control cats were mock-inoculated with Vero E6 cell culture medium supernatant. All animals were maintained individually in Horsfall HEPA-filtered cages. Body temperatures and weight were measured daily until day 7, and then on days 10 and 14 pi. Oropharyngeal (OPS), nasal (NS), and rectal swabs (RS) were collected under sedation (dexmedetomidine) on days 0, 1, 3, 5, 7, 10, and 14 post-inoculation (pi). Upon collection, swabs were placed in sterile tubes containing 1 mL of viral transport medium (VTM Corning®, Glendale, AZ, USA) and stored at −80°C until processed for further analyses. Blood was collected through jugular venipuncture using a 3 mL sterile syringe and 21G × 1” needle and transferred into serum separator tubes on day 4 before inoculation, and on days 0, 7, and 14 pi. The blood tubes were centrifuged at 1200 × g for 10 min and serum was aliquoted and stored at −20°C until further analysis. Three cats from each group were humanely euthanized on days 3, 5, and 14 pi. Following necropsy, bronchoalveolar lavage fluid (BALF), and tissues, including nasal turbinate, palate/tonsil, retropharyngeal LN, trachea, lung, mediastinal LN, heart, liver, spleen, kidney, small intestine, mesenteric LN were collected and processed for rRT-PCR and virus titration. Additionally, nasal turbinate, trachea, and lung were collected and processed for standard microscopic examination and were also processed by *in situ* hybridization (ISH) and *in situ* immunofluorescence (IFA). For this, tissue sections of approximately 0.5 cm in width were fixed by immersion in 10% neutral buffered formalin (20 volumes fixative to 1 volume tissue) for approximately 72 h, and then transferred to 80% ethanol, followed by standard paraffin embedding techniques. Slides for standard microscopic examination were stained with hematoxylin and eosin (HE).

### Nucleic acid isolation and real-time reverse transcriptase PCR (rRT-PCR)

Nucleic acid was extracted from NS, OPS, RS, BALF and tissue samples collected at necropsy. For NS, OPS, RS, and BALF samples 200 µL of cleared swab supernatant were used for nucleic acid extraction. For tissues, 0.3 g of each tissue were minced with a sterile disposable scalpel, resuspended in 3 mL DMEM (10% w/v) and homogenized using a stomacher (one speed cycle of 60s, Stomacher® 80 Biomaster). Then, the tissue homogenate supernatant was centrifuged at 2000 × g for 10 min and 200 µL of cleared supernatant was used for RNA extraction using the MagMax Core extraction kit (Thermo Fisher, Waltham, MA, USA) and the automated KingFisher Flex nucleic acid extractor (Thermo Fisher, Waltham, MA, USA) following the manufacturer’s recommendations. The rRT-PCR for total viral RNA detection was performed using the EZ-SARS-CoV-2 Real-Time RT-PCR assay (Tetracore Inc., Rockville, MD, USA), which detects both genomic and subgenomic viral RNA targeting the virus N gene. An internal inhibition control was included in all reactions. Positive and negative amplification controls were run side-by-side with test samples. For SARS-CoV-2 subgenomic RNA detection, an RT-qPCR reaction targeting the virus envelope (E) gene was used following the primers and protocols previously described [41]. Both RT-PCR (for total viral RNA detection) and RT-qPCR assay (for specific subgenomic RNA detection) were verified using a standard curve by using ten-fold serial dilutions from 10^0^ to 10^−8^ of virus suspension containing 10^6^ TCID_50_.mL^-1^ for each of the SARS-CoV-2 variants used in the study. Relative viral genome copy numbers were calculated based on the standard curve and determined using GraphPad Prism 9 (GraphPad, La Jolla, CA, USA). The amount of viral RNA detected in samples were expressed as log (genome copy number) per mL.

### Virus isolation and titrations

Samples that tested positive for SARS-CoV-2 by rRT-PCR were subjected to virus isolation under Biosafety Level 3 (BSL-3) conditions at the Animal Health Diagnostic Center (ADHC) Research Suite at Cornell University. NS, OPS, RS, BALF, and tissues homogenate supernatant were subjected to end point titrations. For this, samples were subjected to limiting dilutions and inoculated into Vero E6/TMPRSS2 cells cultures prepared 24 hours in advance in 96-well plates. At 48 hours pi, cells were fixed and subjected to IFA as described in a previous study [24]. The limit of detection (LOD) for infectious virus titration is 10^1.05^ TCID_50_.mL^−1^. Virus titers were determined on each time point using end-point dilutions and the Spearman and Karber’s method and expressed as TCID_50_.mL^−1^.

### *In situ* RNA detection

Paraffin-embedded tissues from days 3 and 5 pi were sectioned at 5 µm and subjected to *in situ* hybridization (ISH) using the RNAscope® ZZ probe technology (Advanced Cell Diagnostics, Newark, CA). Nasal turbinate, trachea, and lung from inoculated and controls were subjected to ISH using the RNAscope^®^ 2.5 HD Reagents–RED kit (Advanced Cell Diagnostics) following the manufacturer’s instructions and using a probe targeting SARS-CoV-2 RNA S (V-nCoV2019-S probe ref # 848561). A probe targeting feline host protein peptidylprolyl isomerase B (PPIB) was used as a positive control (Advanced Cell Diagnostics cat # 455011). A probe targeting DapB gene from Bacillus subtilis strain SMY was used as a negative control (Advanced Cell Diagnostics cat # 310043).

### *In situ* immunofluorescence (IFA)

Paraffin-embedded tissues from days 3 and 5 pi were sectioned at 5 µm and subjected to IFA. Tissues from inoculated and control cats including nasal turbinate, palate/tonsil, retropharyngeal LN, trachea, lung, and heart were subjected to IFA. Formalin-fixed paraffin-embedded (FFPE) tissues were deparaffinized with xylene and rehydrated through a series of graded alcohol solutions. Antigen unmasking was performed using Tris-based antigen unmasking solution pH 9.0 (Vector Laboratories ref # H-3301) by boiling the slides in the unmasking solution for 20 minutes. After 10 minutes at 0.2% Triton X-100 (in phosphate-buffered saline [PBS]) at room temperature (RT), and 30 minutes blocking using a goat normal serum (1% in PBS) at RT, tissues were subjected to IFA. A mouse monoclonal antibody targeting SARS-CoV-2 N protein was used as a primary antibody (SARS-CoV-2 N mAb clone B61G11) [24] was incubated for 45 minutes at RT. Followed by 30 minutes incubation at RT with a goat anti-mouse IgG antibody (goat anti-mouse IgG, Alexa Fluor^®^ 488). Nuclear counterstain was performed with 4’,6-Diamidino-2-Phenylindole, Dihydrochloride (DAPI) (10 minutes at RT).

### Histology

For the histological examination, tissue sections of approximately 0.5 cm in width were fixed by immersion in 10% neutral buffered formalin (≥20 volumes fixative to 1 volume tissue) for approximately 72 hours, and then transferred to 80% ethanol, followed by standard paraffin embedding techniques. Nasal turbinate and lung collected from inoculated and control cats were subjected to histological examination after stained with hematoxylin and eosin (HE).

### Luciferase-Immunoprecipitation System (LIPS)

The N protein of SARS-CoV-2 B.1 D614G variant (NYI67-20 strain) was cloned into Nano-Luc vector pNLF1-N [CMV/Hygro] (N terminus) (Cat. N1351, Promega) as a fusion protein. Briefly, The N protein of SARS-CoV-2 was amplified by PCR and cloned in-frame to the 5′end of the NLuc gene between the restriction enzymes *SacII* and *NotI*. A FLAG tag was added to 5′ end of N for the confirmation of expression. The resulting plasmid was sequenced to verify the authenticity of the inserted gene and the expression of the N protein in HEK293T cells was confirmed by IFA and Western blot assays. The NLuc-tagged antigen was generated in HEK293T cells transfected with 10 μg of pNLF1-N plasmid containing SARS-CoV-2 N protein as described previously [26]. At 48 hours post-transfection, cells were lysed with RIPA lysis and extraction buffer (ThermoFisher Scientific, Rockford, IL, USA), supplemented with 1x protease inhibitor (Pierce Protease Inhibitor Tablet, EDTA-Free, ThermoFisher Scientific, Rockford, IL, USA) and the light units (LU) per µl of lysates were measured using Nano-Glo® Luciferase Assay System (Promega) in a luminometer (BioTek Synergy LX Multimode Reader). The LIPS assay was performed following protocols described in a previous study [26] with some modifications. Briefly, 50 µl NLuc-antigen (10^7^ LU per well) and 50 µl serum (heat inactivated and diluted 1:50) were mixed in a 96-well plate and incubated for 1 hour on a rotary shaker at room temperature. Then, the NLuc-antigen and serum mixture were incubated with 10 μl of a 30% suspension of Ultralink protein A/G beads (Pierce Biotechnology, Rockford, IL) in PBS in a 96-well plate. After 1 hour incubation, the antigen-serum-bead mixture was transferred into a 96-well filter HTS plate (Millipore, Bedford, MA). The plate was washed 8 times with Buffer A followed by twice with PBS on a vacuum manifold as described previously [26]. The plate was read using Nano-Glo® Luciferase Assay System (Promega) in a luminometer (BioTek Synergy LX Multimode Reader) and the luminescence (LU) corresponded to the quantity of N protein-specific antibody present in the test serum was calculated. Pooled serum from non-inoculated control cats were used as negative control.

### Neutralizing antibodies

Neutralizing antibody responses to SARS-CoV-2 were assessed by virus neutralization (VN) assay performed under BSL-3 laboratory conditions. Serum samples collected on day 0, 7, and 14 pi were tested against the both recombinant viruses rWA1-D614G and rWA1-Omi-S. For the VN assay, two-fold serial dilutions (1:8 to 1:1,024) of serum samples were incubated with 100 - 200 TCID_50_ of rWA1-D614G or rWA1-Omi-S for 1 hour at 37°C. Following incubation of serum and virus, 50 µL of a cell suspension of Vero E6 cells was added to each well of a 96-well plate and incubated for 48 hours at 37°C with 5% CO_2_. At 48 hours pi, cells were fixed and subjected to IFA as described in a previous study [24]. Neutralizing antibody titers were expressed as the reciprocal of the highest dilution of serum that completely inhibited SARS-CoV-2 infection/replication. FBS and positive and negative serum samples from white-tailed-deer [42] were used as controls.

### Statistical analysis and data plotting

Statistical analysis was performed by 2-way analysis of variance (ANOVA) followed by multiple comparisons. Statistical analysis and data plotting were performed using the GraphPad Prism software (version 9.0.1). Figures 1A and 4A and the graphic abstract were created with BioRender.com.

## Acknowledgments

We thank the Center for Animal Resources and Education (CARE) staff and Cornell Biosafety team for their support. We thank Dr. Cara Mitchel and all staff at Clinvet for their support and donation of the animals.

## Author Contributions

Conceptualization: Diego G. Diel. Data curation: Mathias Martins, Mohammed Nooruzzaman, Jessie Lee Cunningham. Formal analysis: Mathias Martins, Mohammed Nooruzzaman, Jessie Lee Cunningham, Leonardo C. Caserta, Chengjin Ye, Nathaniel Jackson, Diego G. Diel. Funding acquisition: Diego G. Diel. Investigation: Mathias Martins, Mohammed Nooruzzaman, Jessie Lee Cunningham, Leonardo C. Caserta, Diego G. Diel. Methodology: Mathias Martins, Mohammed Nooruzzaman, Jessie Lee Cunningham, Chengjin Ye, Nathaniel Jackson, Leonardo C. Caserta, Luis Martinez-Sobrido, Diego G. Diel. Project administration: Ying Fang, Diego G. Diel. Resources: Luis Martinez-Sobrido, Ying Fang, Diego G. Diel. Supervision: Diego G. Diel. Validation: Mathias Martins, Mohammed Nooruzzaman, Jessie Lee Cunningham. Writing – original draft: Mathias Martins, Mohammed Nooruzzaman, Diego G. Diel. Writing – review & editing: Mathias Martins, Mohammed Nooruzzaman, Chengjin Ye, Luis Martinez-Sobrido, Ying Fang, Diego G. Diel.

## References

1. Zhu N, Zhang D, Wang W, Li X, Yang B, Song J, et al. A novel coronavirus from patients with pneumonia in China, 2019. N Engl J Med. 2020;382: 727–733. doi:10.1056/NEJMoa2001017

2. Worobey M, Levy JI, Malpica Serrano L, Crits-Christoph A, Pekar JE, Goldstein SA, et al. The Huanan Seafood Wholesale Market in Wuhan was the early epicenter of the COVID-19 pandemic. Science. 2022;377: 951–959. doi:10.1126/science.abp8715

3. Holmes EC, Goldstein SA, Rasmussen AL, Robertson DL, Crits-Christoph A, Wertheim JO, et al. The origins of SARS-CoV-2: A critical review. Cell. 2021;184: 4848–4856. doi:10.1016/j.cell.2021.08.017

4. Zhou P, Yang X-L, Wang X-G, Hu B, Zhang L, Zhang W, et al. A pneumonia outbreak associated with a new coronavirus of probable bat origin. Nature. 2020;579: 270–273. doi:10.1038/s41586-020-2012-7

5. Zabidi NZ, Liew HL, Farouk IA, Puniyamurti A, Yip AJW, Wijesinghe VN, et al. Evolution of SARS-CoV-2 variants: Implications on immune escape, vaccination, therapeutic and diagnostic strategies. Viruses. 2023;15: 944. doi:10.3390/v15040944

6. Korber B, Fischer WM, Gnanakaran S, Yoon H, Theiler J, Abfalterer W, et al. Tracking changes in SARS-CoV-2 spike: Evidence that D614G increases infectivity of the COVID-19 virus. Cell. 2020;182: 812–827.e19. doi:10.1016/j.cell.2020.06.043

7. World Health Organization. Tracking SARS-CoV-2 variants. Geneva, Switzerland; 2023.

8. World Health Organization. Classification of Omicron (B.1.1.529): SARS-CoV-2 Variant of Concern. Geneva, Switzerland; 2021 Nov.

9. Iuliano AD, Brunkard JM, Boehmer TK, Peterson E, Adjei S, Binder AM, et al. Trends in disease severity and health care utilization during the early Omicron variant period compared with previous SARS-CoV-2 high transmission periods — United States, December 2020–January 2022. MMWR Morb Mortal Wkly Rep. 2022;71: 146–152. doi:10.15585/mmwr.mm7104e4

10. Dejnirattisai W, Shaw RH, Supasa P, Liu C, Stuart AS, Pollard AJ, et al. Reduced neutralisation of SARS-CoV-2 omicron B.1.1.529 variant by post-immunisation serum. Lancet. 2022;399: 234–236. doi:10.1016/S0140-6736(21)02844-0

11. Ai J, Zhang H, Zhang Y, Lin K, Zhang Y, Wu J, et al. Omicron variant showed lower neutralizing sensitivity than other SARS-CoV-2 variants to immune sera elicited by vaccines after boost. Emerg Microbes Infect. 2022;11: 337–343. doi:10.1080/22221751.2021.2022440

12. Zhang L, Li Q, Liang Z, Li T, Liu S, Cui Q, et al. The significant immune escape of pseudotyped SARS-CoV-2 variant Omicron. Emerg Microbes Infect. 2022;11: 1–5. doi:10.1080/22221751.2021.2017757

13. Cele S, Jackson L, Khoury DS, Khan K, Moyo-Gwete T, Tegally H, et al. Omicron extensively but incompletely escapes Pfizer BNT162b2 neutralization. Nature. 2022;602: 654–656. doi:10.1038/s41586-021-04387-1

14. Dejnirattisai W, Huo J, Zhou D, Zahradník J, Supasa P, Liu C, et al. SARS-CoV-2 Omicron-B.1.1.529 leads to widespread escape from neutralizing antibody responses. Cell. 2022;185: 467–484.e15. doi:10.1016/j.cell.2021.12.046

15. Schmidt F, Muecksch F, Weisblum Y, Da Silva J, Bednarski E, Cho A, et al. Plasma Neutralization of the SARS-CoV-2 Omicron Variant. N Engl J Med. 2022;386: 599–601. doi:10.1056/NEJMc2119641

16. Mohseni Afshar Z, Tavakoli Pirzaman A, Karim B, Rahimipour Anaraki S, Hosseinzadeh R, Sanjari Pireivatlou E, et al. SARS-CoV-2 Omicron (B.1.1.529) variant: A challenge with COVID-19. Diagnostics. 2023;13: 559. doi:10.3390/diagnostics13030559

17. Chen D-Y, Chin CV, Kenney D, Tavares AH, Khan N, Conway HL, et al. Spike and nsp6 are key determinants of SARS-CoV-2 Omicron BA.1 attenuation. Nature. 2023;615: 143–150. doi:10.1038/s41586-023-05697-2

18. Meng B, Abdullahi A, Ferreira IATM, Goonawardane N, Saito A, Kimura I, et al. Altered TMPRSS2 usage by SARS-CoV-2 Omicron impacts infectivity and fusogenicity. Nature. 2022;603: 706–714. doi:10.1038/s41586-022-04474-x

19. Qu P, Evans JP, Kurhade C, Zeng C, Zheng Y-M, Xu K, et al. Determinants and mechanisms of the low fusogenicity and high dependence on endosomal entry of Omicron subvariants. mBio. 2023;14. doi:10.1128/mbio.03176-22

20. Hu B, Chan JF-W, Liu H, Liu Y, Chai Y, Shi J, et al. Spike mutations contributing to the altered entry preference of SARS-CoV-2 omicron BA.1 and BA.2. Emerg Microbes Infect. 2022;11: 2275–2287. doi:10.1080/22221751.2022.2117098

21. Hu B, Chan JF-W, Liu H, Liu Y, Chai Y, Shi J, et al. Spike mutations contributing to the altered entry preference of SARS-CoV-2 omicron BA.1 and BA.2. Emerg Microbes Infect. 2022;11: 2275–2287. doi:10.1080/22221751.2022.2117098

22. Winkler ES, Chen RE, Alam F, Yildiz S, Case JB, Uccellini MB, et al. SARS-CoV-2 Causes Lung infection without severe disease in human ACE2 knock-in mice. J Virol. 2022;96. doi:10.1128/JVI.01511-21

23. Halfmann PJ, Iida S, Iwatsuki-Horimoto K, Maemura T, Kiso M, Scheaffer SM, et al. SARS-CoV-2 Omicron virus causes attenuated disease in mice and hamsters. Nature. 2022;603: 687–692. doi:10.1038/s41586-022-04441-6

24. Martins M, do Nascimento GM, Nooruzzaman M, Yuan F, Chen C, Caserta LC, et al. The Omicron variant BA.1.1 presents a lower pathogenicity than B.1 D614G and Delta variants in a feline model of SARS-CoV-2 infection. J Virol. 2022;96. doi:10.1128/jvi.00961-22

25. Freitas RS, Crum TF, Parvatiyar K. SARS-CoV-2 Spike Antagonizes innate antiviral immunity by targeting interferon regulatory factor 3. Front Cell Infect Microbiol. 2022;11. doi:10.3389/fcimb.2021.789462

26. Burbelo PD, Ching KH, Klimavicz CM, Iadarola MJ. Antibody profiling by luciferase immunoprecipitation systems (LIPS). J Vis Exp. 2009. doi:10.3791/1549

27. Ren W, Zhang Y, Rao J, Wang Z, Lan J, Liu K, et al. Evolution of immune evasion and host range expansion by the SARS-CoV-2 B.1.1.529 (Omicron) variant. mBio. 2023;14. doi:10.1128/mbio.00416-23

28. Neerukonda SN, Wang R, Vassell R, Baha H, Lusvarghi S, Liu S, et al. Characterization of entry pathways, species-specific angiotensin-converting enzyme 2 residues determining entry, and antibody neutralization evasion of Omicron BA.1, BA.1.1, BA.2, and BA.3 variants. J Virol. 2022;96. doi:10.1128/jvi.01140-22

29. Kim S, Liu Y, Ziarnik M, Seo S, Cao Y, Zhang XF, et al. Binding of human ACE2 and RBD of Omicron enhanced by unique interaction patterns among SARS-CoV-2 variants of concern. J Comput Chem. 2023;44: 594–601. doi:10.1002/jcc.27025

30. McCallum M, Czudnochowski N, Rosen LE, Zepeda SK, Bowen JE, Walls AC, et al. Structural basis of SARS-CoV-2 Omicron immune evasion and receptor engagement. Science. 2022;375: 864–868. doi:10.1126/science.abn8652

31. Suzuki R, Yamasoba D, Kimura I, Wang L, Kishimoto M, Ito J, et al. Attenuated fusogenicity and pathogenicity of SARS-CoV-2 Omicron variant. Nature. 2022;603: 700–705. doi:10.1038/s41586-022-04462-1

32. Qu P, Evans JP, Kurhade C, Zeng C, Zheng Y-M, Xu K, et al. Determinants and mechanisms of the low fusogenicity and high dependence on endosomal entry of Omicron subvariants. mBio. 2023;14. doi:10.1128/mbio.03176-22

33. Zhang Q, Chen Z, Huang C, Sun J, Xue M, Feng T, et al. Severe acute respiratory syndrome coronavirus 2 (SARS-CoV-2) membrane (M) and spike (S) proteins antagonize host type I interferon response. Front Cell Infect Microbiol. 2021;11. doi:10.3389/fcimb.2021.766922

34. Shuai H, Chan JF-W, Hu B, Chai Y, Yuen TT-T, Yin F, et al. Attenuated replication and pathogenicity of SARS-CoV-2 B.1.1.529 Omicron. Nature. 2022;603: 693–699. doi:10.1038/s41586-022-04442-5

35. Yuan S, Ye Z-W, Liang R, Tang K, Zhang AJ, Lu G, et al. Pathogenicity, transmissibility, and fitness of SARS-CoV-2 Omicron in Syrian hamsters. Science. 2022;377: 428–433. doi:10.1126/science.abn8939

36. Nooruzzaman M, Diel DG. Infection dynamics, pathogenesis, and immunity to SARS-CoV-2 in naturally susceptible animal species. J Immunol. 2023;211: 1195–1201. doi:10.4049/jimmunol.2300378

37. Su W, Choy KT, Gu H, Sia SF, Cheng KM, Nizami SIN, et al. Reduced pathogenicity and transmission potential of Omicron BA.1 and BA.2 sublineages compared with the early severe acute respiratory syndrome coronavirus 2 D614G variant in Syrian hamsters. J Infect Dis. 2023;227: 1143–1152. doi:10.1093/infdis/jiac276

38. Liu S, Selvaraj P, Sangare K, Luan B, Wang TT. Spike protein-independent attenuation of SARS-CoV-2 Omicron variant in laboratory mice. Cell Rep. 2022;40: 111359. doi:10.1016/j.celrep.2022.111359

39. Imbiakha B, Ezzatpour S, Buchholz DW, Sahler J, Ye C, Olarte-Castillo XA, et al. Age-dependent acquisition of pathogenicity by SARS-CoV-2 Omicron BA.5. Sci Adv. 2023;9: eadj1736. doi:10.1126/sciadv.adj1736

40. Ye C, Chiem K, Park J-G, Oladunni F, Platt RN, Anderson T, et al. Rescue of SARS-CoV-2 from a Single Bacterial Artificial Chromosome. mBio. 2020;11. doi:10.1128/mBio.02168-20

41. Dagotto G, Mercado NB, Martinez DR, Hou YJ, Nkolola JP, Carnahan RH, et al. Comparison of subgenomic and total RNA in SARS-CoV-2-challenged Rhesus macaques. J Virol. 2021;95. doi:10.1128/JVI.02370-20

42. Palmer M V., Martins M, Falkenberg S, Buckley A, Caserta LC, Mitchell PK, et al. Susceptibility of white-tailed deer (*Odocoileus virginianus*) to SARS-CoV-2. J Virol. 2021;95. doi:10.1128/JVI.00083-21

